# Behavioral divergence across replicated predation regimes is species-specific in two sympatric poeciliid fishes

**DOI:** 10.64898/2026.10.26.761983

**Authors:** Callen M. Inman, Mahita S. Velaga, Alejandro Gutierrez, Molly E. Cummings

## Abstract

Predation is a potent driver of behavioral divergence across prey populations, yet whether shared predation regimes generate similar responses in coexisting prey species remains unclear. Species differences in reproductive and foraging traits may alter vulnerability to predators, generating species-specific responses. We compared behavior and cognition of two poeciliid fishes (*Gambusia affinis* and *Poecilia latipinna*) that differ in body size, the presence of male ornamentation, and mating tactics. We sampled both species across replicate high- and low-predation regimes and measured exploration, shyness, boldness, social motivation, and detour solve latency (N = 509). Multivariate behavior and cognition differed between predation regimes, and the magnitude of divergence varied with species but not body size. Consistent with the multivariate pattern, the more conspicuous *P. latipinna* exhibited higher boldness and overall lower exploration under high-predation, while high- and low-predation populations of *G. affinis* did not differ. These findings suggest either that the more conspicuous species experiences greater predation risk or that the less conspicuous species reduces its predation risk through alternative phenotypic responses. More broadly, species differences in reproductive phenotypes associated with sexual selection may help explain why shared predation regimes generate variable patterns of behavioral divergence across taxa.

## Introduction

Predation can drive predictable behavioral diversification (Dayton et al. 2005; Eroukhmanoff and Svensson 2009; Archard et al. 2012; Wund et al. 2015; Heinen-Kay et al. 2016; Jiang and Mikolajewski 2018; Langerhans 2018; Lapiedra et al. 2018). However, whether shared predation regimes generate similar behavioral divergence across coexisting prey species remains unclear.

Theory predicts that prey sharing predators need not have identical responses because predator-mediated selection depends on species ecology and phenotype (Abrams 2000). Empirical studies similarly demonstrate that behavioral responses to predation may differ among species exposed to identical predator communities (Chai and Srygley 1990; Randall et al. 1995; Relyea 2001; Richardson 2001; Wirsing et al. 2010; Palmer and Packer 2021). Species-specific responses are expected since prey traits can differentially impact the risk of predation from encounter and detection to pursuit and capture (Endler 1991; Vamosi 2005; Langerhans 2007). For instance, prey that are more detectable, more frequently encountered, or more profitable to predators may compensate for higher predation risk by showing more pronounced anti-predator behavior, such as longer sheltering (Cabido et al. 2009), faster fleeing (Martín et al. 2009), or reduced movement that facilitates crypsis (Relyea 2001; Richardson 2001). Consequently, even when sympatric prey species experience identical predator communities, differences in morphology, habitat use, and reproductive conspicuousness may cause identical predation regimes to generate unique patterns of behavioral divergence.

One important class of traits that may alter predation risk comprises reproductive phenotypes shaped by sexual selection (Haskins et al. 1961; Magnhagen 1991; Sih 1994). Conspicuous display morphology and behavior can increase predation risk (Winemiller et al. 1990; Rosenthal et al. 2001; Basolo and Wagner 2004; Pruden and Uetz 2004; Fowler-Finn and Hebets 2011; Clark et al. 2018; Whitaker et al. 2021), leading to spatial or temporal shifts in reproductive behavior (Endler 1987; Magurran and Seghers 1990; Fuller and Berglund 1996). Co-occurring species that differ in the conspicuousness of their ornamentation and mating behavior may experience different predator-mediated tradeoffs, and this cascading effect could potentially lead to differences in behaviors outside the reproductive context, including risk-taking, exploration, and cognitive flexibility. Reproductive phenotypes may also be associated with behavioral and cognitive differences independently of their effects on predation risk (Dadda 2015; Cummings et al. 2022; Queller et al. 2022; Queller et al. 2023; Queller et al. 2024; Inman et al. 2025a). However, whether such developmental or species differences in reproductive phenotypes modify behavioral responses across predation regimes is poorly understood.

In addition to species differences in reproductive phenotype, individual variation in body size may independently alter predation risk. Although prey size may limit successful consumption by gape-limited predators (Mihalitsis and Bellwood 2017), larger prey are more detectable (Eriksson 1985) and provide greater energetic rewards (Krebs et al. 1977), leading many predators to preferentially attack larger individuals (Britton and Moser 1982; Trexler et al. 1994; Tobler et al. 2007; Johansson et al. 2004). As a result, larger individuals may experience different tradeoffs between risk avoidance and activities such as foraging or reproduction, potentially favoring different optimal behavioral phenotypes. Accordingly, behaviors such as boldness and exploration have been found to be size-specific (Brown and Braithwaite 2004; Brown et al. 2007b; Kelleher et al. 2017). Such differences may reflect adaptive responses to size-specific predation risk, energetic requirements, or locomotor constraints that covary with body size (Clarke and Johnston 1999). In turn, body size may modify behavioral responses to predation independently of species identity.

Previous work on species-specific responses to predation has largely focused on direct anti-predator behavior. However, predation environments may shape behaviors and cognitive abilities expressed outside of a predator-prey encounter such as boldness, exploration, and cognitive flexibility (Réale et al. 2007; Sih et al. 2012; Heinen-Kay et al. 2016; Vila Pouca et al. 2021). These traits may influence how individuals balance predator avoidance against other activities such as foraging and mating, and therefore may also shift in response to persistent differences in predation risk (Lima and Dill 1990; Werner and Anholt 1993; Réale and Festa-Bianchet 2003; Koch et al. 2007; Hay et al. 2008; Ballew et al. 2017). Levels of boldness and exploration are often correlated with one another across individuals and vary by predation regime (Archard and Braithwaite 2011; Harris et al. 2010; Money et al. 2017). This suggests that predation might influence a suite of behaviors and cognitive abilities outside of the direct predator-prey interaction. Additionally, cognitive flexibility may facilitate rapid behavioral adjustments in response to predation risk. Constant vigilance can be costly to activities such as foraging and mating (Metcalfe 1984; Lima 1987). Individuals capable of rapidly modifying their behavior when predators are encountered thus may allocate less time to vigilance (Lima and Dill 1990; Herberholz and Marquart 2012). Consistent with this hypothesis, research in guppies has demonstrated that early-life experience with simulated predation improves cognitive flexibility in adult guppies (Vila Pouca et al. 2021), although comparable studies remain scarce.

Empirical studies of boldness and exploration across different levels of predation threat are more numerous but have produced inconsistent results. For example, boldness favors survival of prey fish against certain predators (Blake et al. 2018, Fuiman et al. 2010) but may lower survival against others (Hulthén et al. 2017). Accordingly, interpopulation studies of boldness between predation regimes have found opposing patterns across fish taxa (Brown et al. 2005; Brown et al. 2007a; Harris et al. 2010; Heinen-Kay et al. 2016). Such inconsistencies may arise because species differ in the phenotypic traits that mediate predator-prey interactions.

Here, we examine how predation risk shapes behavior and cognition in two sympatric poeciliid fishes, *Gambusia affinis* and *Poecilia latipinna*. Poeciliids display dramatic interspecific variation in reproductive behavior and morphology, ranging from species in which drab males rely exclusively on coercive sneak copulations to species in which ornamented males perform elaborate courtship displays (Furness et al. 2019; Reznick et al. 2021). Many poeciliid species also occupy environments that differ in predator abundance (Meffe and Snelson 1989) and exhibit predator-associated divergence in behavior and reproductive traits (Haskins et al. 1961; Seghers 1974; Endler 1978; Endler 1980; Magurran and Seghers 1990; Brown and Braithwaite 2004; Brown et al. 2005; Harris et al. 2010; Heinen-Kay et al. 2016). Male mating tactics in *G. affinis* are dominated by inconspicuous coercive sneak copulations (Bisazza et al. 2001; Deaton 2008). Meanwhile, *P. latipinna* males exhibit alternative mating tactics: small males primarily coerce, while large males with conspicuous dorsal ‘sail’ fins and lateral body coloration primarily court females (Baird 1974; Farr et al. 1986). Because these two species with differing levels of conspicuousness co-occur across replicate environments that vary markedly in predator abundance, they provide an opportunity for examining whether sympatric species show shared or species-specific behavioral divergence across replicated predation regimes.

Using both male and female wild fish collected from replicate pairs of high- and low-predation locations, we quantified exploration, boldness and shyness, social motivation, and inhibitory control, a type of cognitive flexibility. We first ask whether the two sympatric species exhibited shared or species-specific patterns of multivariate behavioral divergence across predation regimes. We then ask whether predation-associated divergence in individual behavioral and cognition is (1) consistent across replicated predation regimes, (2) stronger in the more conspicuous *P. latipinna*, and (3) stronger among larger-bodied individuals.

## Methods

### Field collection

509 sexually mature fish were collected from paired field sites in three major central Texas watersheds (Brushy Creek, San Marcos River, and Guadalupe River), representing replicate high- and low-predation regimes (Figure 1) between February 2024 and July 2025. Fish were collected with Scientific Research Permits (2023-2025) from the Texas Parks & Wildlife Department and permission of UT Austin IACUC. Fish were collected as early in the year as February and as late in the year as October to coincide with the known breeding seasons of *P. latipinna* and *G. affinis* at similar latitudes (Baird 1974; Self 1940; Medlen 1951). The two species were collected using large dip nets at each site. At each site, *G. affinis* and *P. latipinna* occupied similar microhabitats in mixed-species shoals. Field sites were identified based on the presence or absence of piscivorous fish. Because avian predators are highly mobile and patchy in nature compared to piscine predators (e.g. Kloskowski 2011), we did not use their presence or absence as a criterion for determining whether a site was high- or low-predation. However, avian predators tended to be absent at sites without piscine predators and present at sites with piscine predators (Inman, C.M., personal observation). Besides the fact that high-predation sites tended to have deeper water and higher flow than low-predation sites, many other environmental attributes likely varied among sites, but not with respect to predation regime (e.g. structural complexity, water temperature) (Figure 1).

**Figure 1.**
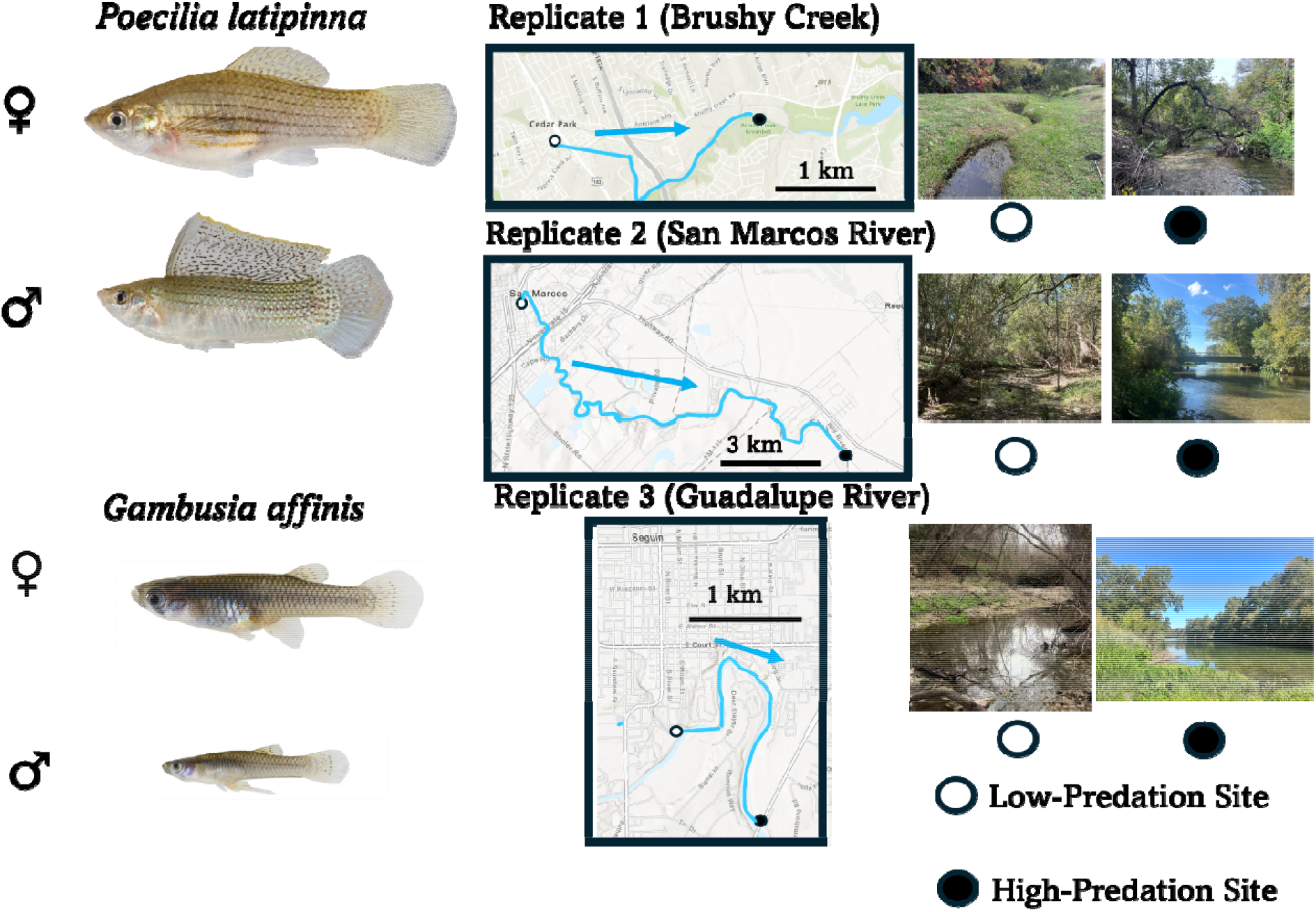
Left: Representative photographs of adult *Poecilia latipinna* (top) and *Gambusia affinis* (bottom) of each sex. Center: Maps of the 3 replicate watersheds in central Texas. Blue arrows indicate the direction of flow: high-predation sites were located downstream of low-predation sites. Right: Photos of each paired low- and high-predation field site. Open circles indicate low-predation sites, while closed circles indicate high-predation sites. High-predation sites had steep depth gradients (2-6 m maximum depth), whereas low-predation sites were consistently shallow (<1 m max. depth) pools or slow-flowing streams.

### Replicate 1: Brushy Creek

In the Brushy Creek drainage, the low-predation site (Brushy Creek Headwaters: 30.504133876974716, -97.82013337761556) consisted of two isolated pools containing *Poecilia latipinna* and *Gambusia affinis* and no other fish species. At the high-predation site 2.5 km downstream of the low-predation site (Brushy Creek Lake Park: 30.505011785075904, - 97.78619363679086), observers (CI, MH, AV, MR) noted the presence of adult green sunfish (*Lepomis cyanellus*), bluegill (*Lepomis macrochirus*), longear sunfish (*Lepomis megalotis*), Rio Grande cichlid (*Herichthys cyanoguttatus*), and largemouth bass (*Micropterus nigricans*), with between 10-15 individuals observed on any one visit. 98 fish were collected from the Brushy Creek low-predation site (*G. affinis*: 29 females, 26 males; *P. latipinna*: 17 females, 20 males), and 92 fish were collected from the Brushy Creek high-predation site (*G. affinis*: 22 females, 13 males; *P. latipinna*: 20 females, 11 males). Individuals from both locations and species were transported to J.T. Patterson Labs at The University of Texas at Austin and assayed between February 2024 and July 2024 and in July 2025.

### Replicate 2: San Marcos River

In the San Marcos River watershed, the low-predation site (Purgatory Creek: 29.881157111633925, -97.93641354354374) contained *G. affinis* and *P. latipinna* as the two dominant fish species. No piscine predators were observed during multiple site visits between September 2024 and August 2025. At the high-predation site 15.4 km downstream of the low-predation site (Martindale: 29.849571508531245, -97.8569867870611), the two focal species co-occurred with several piscine predators along a steep depth gradient. During site visits, black basses (*Micropterus* sp.) and Rio Grande cichlid were the main predators recorded, with a typical abundance of 10-20 individual predators. Research grade iNaturalist records taken at the location during the sampling period also record redear sunfish (*Lepomis microlophus*) and channel catfish (*Ictalurus punctatus*). 83 fish were collected from the San Marcos River watershed low-predation site (*G. affinis*: 21 females, 20 males; *P. latipinna*: 25 females, 17 males) and 100 fish from the San Marcos River watershed high-predation site (*G. affinis*: 21 females, 22 males; *P. latipinna*: 25 females, 32 males). Individuals from both locations and species were transported to J.T. Patterson Labs for behavioral experiments between March and July 2025.

### Replicate 3: Guadalupe River

Finally, in the Guadalupe River drainage, the low-predation site (Seguin-Walnut Branch: 29.559379408056124, -97.96163102206515) contained only *P. latipinna* and *G. affinis* in a series of shallow pools. No piscine predators were observed during the sampling period, from March to August of 2025. The high-predation Guadalupe River site (Seguin-RV Park: 29.546501283255235, -97.95028720821772) is 3.80 km downstream of the lower-predation site on the main stem of the Guadalupe River. Here, we observed black basses, sunfishes (*Lepomis* sp.), and Rio Grande cichlid during the sampling period. 84 fish were collected from the Guadalupe River low-predation site (*G. affinis*: 22 females, 22 males; *P. latipinna*: 25 females, 15 males), and 84 fish were also collected from the Guadalupe River high-predation site (*G. affinis*: 26 females, 12 males; *P. latipinna*: 28 females, 18 males). Individuals of both species were collected, transported to J.T. Patterson Labs, and assayed between March 2025 and August 2025.

### Laboratory Housing and Behavioral Assays

Fish were added to 75-L species- and site-specific community tanks with gravel bottoms and synthetic plants in the laboratory at UT Austin, where they were acclimated for 5-12 days before behavioral and cognitive assays began. Fish were fed on a daily basis with a mixture of TetraMin flakes and Aquaxcel starter fish pellets. Tanks were aerated and maintained at a temperature of 24°C on a 12-hour day: night cycle. Fish husbandry and experiments were approved by the UT Austin Institutional Animal Care and Use Committee (protocol number AUP-2025-2200224).

Once the acclimation period ended, individuals were then socially isolated for a 2-day period prior to the behavioral and cognitive assays. Fish could interact with individuals in neighboring isolation tanks (via a shared transparent tank wall), but only one individual was present in each tank. To reduce aggression on either side of the tank barrier, males were never isolated adjacent to other males. Following the acclimation period, we tested individual fish in a detour reaching task and a scototaxis assay (Figure 2). To limit the potential carryover effects of assay stress (Bell 2013), fish completed detour reaching before scototaxis, with three days in social isolation between assays. Once assays were completed, we photographed each individual fish under standardized conditions to collect morphometric data (including standard length), which was later processed using ImageJ (Schneider et al. 2012).

**Figure 2.**
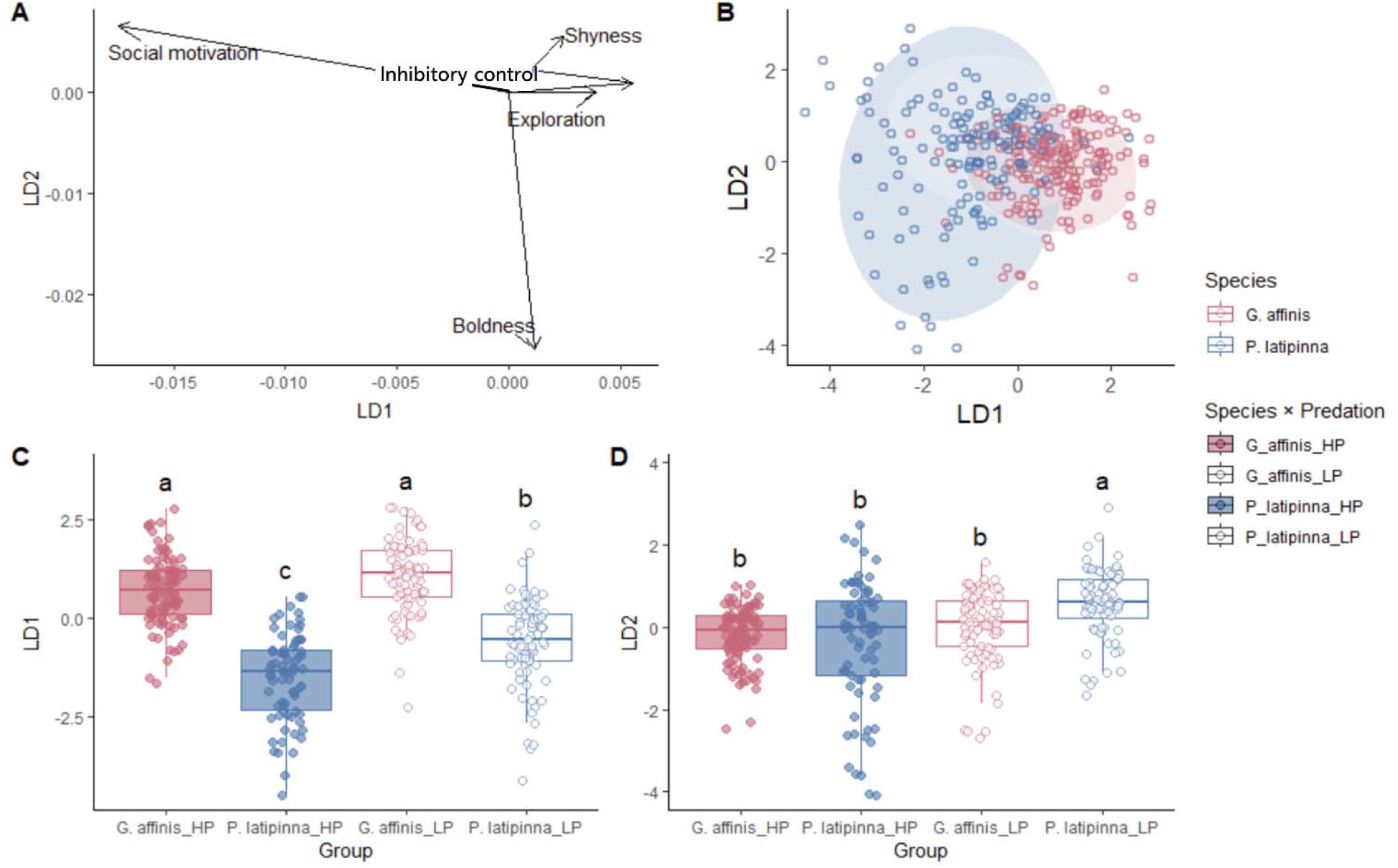
Results from Linear Discriminant Analysis that grouped individuals by species and predation environment (HP and LP). Panel (A) shows loadings onto the two major axes differentiating the four groups. Panel (B) plots individuals onto 95% confidence interval ellipses based on species and predation environment. Open points indicate low-predation, while filled points indicate high-predation. Panel (C) shows variation along LD1 axis. Letters indicate significant (p < 0.05) differences according to post-hoc Tukey tests performed on ANOVA results. Panel (D) shows variation along the LD2 axis.

### Detour reaching

We used a detour reaching task to measure both social motivation and inhibitory control (Inman et al. 2025b), defined as an organism’s ability to inhibit a prepotent response to a stimulus according to new information (Kabadayi et al. 2018; van Horik et al. 2018). We began the assay by placing a conspecific female reward fish in a transparent glass cylinder on the opposite side of the transparent barrier from the starting point (Inman et al. 2025b). The focal fish underwent a 5-minute habituation period behind an opaque barrier before starting the trial. In each trial, the barrier was removed, and the focal fish was recorded for a 10-minute observation period using a Nikon D3300 camera. Detour video recordings were hand-scored by an undergraduate student (MV), unaware of the objectives of the study, for two measures: latency to engage with glass barrier and solve time. We measured latency to engage in the task by calculating the time it took each focal fish to travel from the start position (release from habituation) to the glass barrier (within 2 cm; see Figure 3) or to turn its body fully parallel to the glass barrier after exiting the habituation alley. Because the reward fish was visible to the focal individual from the beginning of the trial, we considered this engagement latency a measure of social motivation. We measured detour solve time, a measure of inhibitory control, as the time it took each fish to travel from the glass barrier to the reward zone (defined as more than half of its body passing through the reward zone entryway).

**Figure 3.**
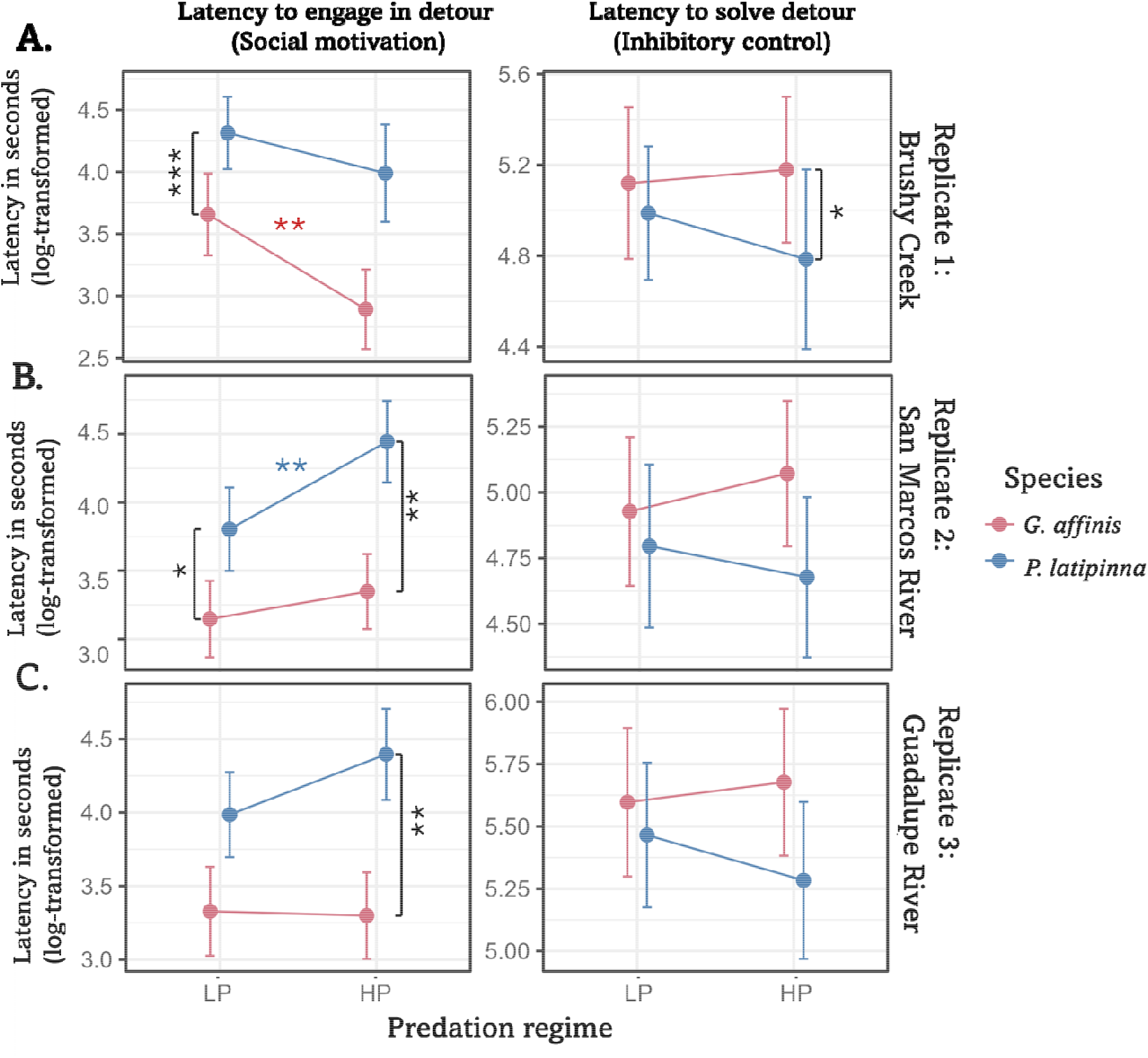
Estimates of social motivation (social latency) and inhibitory control(problem solving latency) measures by species and predation regime in (A.) Replicate 1 (Brushy Creek), (B.) Replicate 2 (San Marcos River), and (C.) Replicate 3 (Guadalupe River). Vertical asterisks and square brackets indicate significant species differences in model-adjusted estimated marginal means at a particular location. Horizontal asterisks are color-coded by species and indicate significant differences in estimated marginal means between predation regimes within a watershed. For a given contrast, one asterisk indicates a p-value < 0.05, two indicates a p-value < 0.01, and three indicates a p-value < 0.001. Estimated marginal means for detecting species differences are derived from linear models (LMs) that include a three-way interaction between species, predation regime, and watershed, in addition to sex and standard length covariates. Estimated marginal means for predation regime differences use the same LMs employed elsewhere in the study (see Methods).

### Scototaxis

The scototaxis assay simulates exposed and sheltered areas through a tank divided into bright (white) and dark (black halves), respectively. This assay has been used to assess shyness–boldness attributes in a number of small fish species. These species, including some poeciliids, have consistently shown aversive responses to the white zone and tend to spend more time in the black zone (Maximino et al. 2010). To begin each trial, we placed each fish in the middle of the tank between a black barrier and a white barrier for a 5-minute habituation period. Following habituation, we removed the barriers and began video recording for a 10 min period. After each individual trial, we completed a full water change of the scototaxis tank. We used Ethovision XT15 (Noldus et al. 2001) to quantify three behaviors: shyness, boldness, and exploratory behavior. To assess boldness, we measured the amount of time each individual spent in the white half of the tank (‘time in white’). To assess shyness, we measured the amount of time each individual spent in black thigmotaxis, or the 2-cm zone on the edge of the black zone closest to the tank wall (see Inman et al. 2025b). We used total distance moved (cm) to capture exploration. This metric differs from a more general activity measure collected in a non-threatening context (Dingemanse et al. 2007; Réale et al. 2007; Etheredge et al. 2018; Inman et al. 2025a). A subset of individuals (n = 101) underwent the scototaxis assay twice, with an interval of four days in between trials, to assess individual-level repeatability. 52.5% of the sample (n = 53) were *G. affinis*, while 47.5% (n = 48) were *P. latipinna*. 29.7% of the sample (n = 30) were from high-predation regimes, while 70.3% of the sample (n = 71) were from low-predation regimes. 95% of the retested individuals (n = 96) were from the Brushy Creek watershed.

### Statistical analysis

Statistical analyses were conducted using R version 4.6.0 (R Core Team 2025). First, we used Principal Component Analysis to verify that the scototaxis variables of interest (total distance moved (exploration), time in the white zone (boldness), and time in black thigmotaxis (shyness)) mapped onto independent axes and were not redundant variables. A broken-stick model was used to determine whether the variance explained by each PC was larger than the null expectation. If more variance is explained than the null expectation by the first two principal components, this challenges the idea that time in white, time in black thigmotaxis, and total distance moved are independent axes.

Next, we constructed a MANCOVA model to test whether multivariate behavior and cognition varied with predation regime and whether the association differed by replicate, species, or body size. The focal behaviors and cognitive abilities in which we were interested, described in the paragraphs above, were problem-solving latency in detour, latency to barrier engagement in detour, distance moved in scototaxis, time in scototaxis white zone, and time in black thigmotaxis. After log-transforming each behavioral or cognitive response to fit model assumptions (Huberty and Petoskey 2000) and scaling standard length, we constructed a MANCOVA model with the five behaviors as response variables. Due to the presence of some “0” values in the dataset, “0.01” was added to all “time in black thigmotaxis” and “time in white zone” values prior to log-transformation.

We included predation regime as a main effect as well as a replicate watershed * predation regime interaction term (Arendt et al. 2025). We also included predation regime * species and predation regime * standard length interaction terms. Standard length and sex were included as covariates. We estimated effect sizes for fixed effects in the MANCOVA model using F-values and partial η² computed from Type III F tests for Wilks’ Lambda using the R package ‘car’ (Fox and Weisberg 2019). Partial η² describes the proportion of variation explained by each independent factor in a (M)ANOVA model (Arendt et al. 2025). Partial η² effect sizes were interpreted according to Cohen (1969).

We also conducted a second MANCOVA that included an interaction term among sex, species, and predation regime to confirm that the hypothesized species * predation regime was not highly sex-dependent, as has been observed in other studies (Godin 1995; Brusseau et al. 2023).

To assess how species and predation regime combinations differed in suites of behaviors, we performed a Linear Discriminant Analysis (LDA). We used four pre-defined categories of *G. affinis* High-Predation, *G. affinis* Low-Predation, *P. latipinna* High-Predation, and *P. latipinna* Low-Predation based on the five focal behaviors (social latency, solve latency, exploration, boldness, and shyness). We performed leave-one-out cross-validation (LOOCV) on the LDA model to estimate the percentage of individuals that were correctly assigned to each of these categories.

Linear models (LMs) used the same focal interaction terms as the MANCOVA to test replicate-, species- and size-dependent associations with predation regime. As we did for the MANCOVA, we also constructed a separate set of models that included sex as an interaction term. We initially included population as a random intercept in linear mixed-effects models; however, all models estimated the population-level variance as zero and produced singular fits. Because the random effect contributed no estimated variance, we report the simpler LMs instead. We evaluated model terms using Type III F-tests and calculated partial η² as an effect-size measure. To account for multiple testing across responses, p-values for each model term were adjusted across the five response variables using the false discovery rate procedure (Benjamini and Hochberg 1995; Pike 2011).

After performing each LM, we used assessed residual uniformity, dispersion, and outliers using conventional and DHARMa diagnostics (Hartig 2024; Table S6). Models with detour response variables did not have substantial violations (Figure S2; Table S6), whereas scototaxis models showed lower-tail deviations associated with zero or near-zero responses (Figure S3; Table S6). Sensitivity analyses excluding observations with standardized residuals > 3 did not alter focal Species * Predation Regime or Standard Length * Predation Regime results (Table S7). We therefore retained the original linear models (Table S7).

We then used estimated marginal means (EMMs), calculated with the R package “emmeans” (Lenth 2025) to characterize the direction and magnitude of significant effects and interactions. In particular, we examined differences between high- and low-predation populations within species and replicate watersheds to assess the direction and consistency of predation regime associations shown by omnibus models.

To estimate individual-level repeatability for each of the three focal behaviors in the scototaxis assay, we used the “rptR” package (Stoffel et al. 2017) to construct linear mixed models (LMMs) with each log-transformed behavioral measure as the response, Trial number (1 or 2) as the fixed effect, and individual identity as a random intercept. We estimated as the proportion of total variance attributable to individual differences using intraclass correlation coefficients (Bell et al. 2009). We used a parametric bootstrap of 1000 replications to calculate a 95% confidence interval of repeatability for each variable. We also implemented a randomization procedure of 1000 permutations on residuals of the null model to carry out likelihood ratio tests.

## Results

We assessed behavior, cognition, and morphometrics in a total of 509 individuals: 253 *P. latipinna* (140 females and 113 males) and 256 *G. affinis* (141 females and 115 males).

All three scototaxis variables were significantly repeatable in the subset of individuals we tested twice (n = 101; p < 0.01, permutation test; Table S5). Total distance moved was the most repeatable behavior (R = 0.619, 95% CI = 0.481-0.724, p = 0.001; Table S5), while time in white (R = 0.389, 95% CI = 0.218-0.537; Table S5) and time in black thigmotaxis (R = 0.353, CI = 0.18-0.511; Table S5) both had moderate repeatability. Predation regime and species-specific estimates are presented descriptively in the Supplementary Material (Table S5).

In the Principal Component Analysis conducted on the three focal variables in scototaxis (time in white, time in black thigmotaxis, distance moved), we found that all three variables were similar in PC1 (51.34% of the variance), but that the three variables differed from one another in PC2 (25.81% of the variance), and PC3 (22.83% of the variance). The null expectation for the variance explained by the first principal component is 0.611, the second component 0.278, and the third component 0.111. Thus, PC1 and PC2 explain less of the variance than expected, while PC3 explains more of the variance than expected at random. This indicates that time in white, time in black thigmotaxis, and distance moved are indeed separable axes that do not show a strong common axis (PC1) or secondary structure (PC2). We did not find meaningful differences between the two MANCOVAs we conducted or significant interactions among sex, species, and predation regime in the MANCOVA that included sex dependencies. Thus, we only report the MANCOVA that included a three-way interaction term with sex in the Supplementary Material (Table S3). In the primary MANCOVA, we found that predation regime had a large effect on behavioral and cognitive variation and covariation (partial ^2^ = 0.15, p < 0.001; Table 1). The interaction between predation regime and replicate had a smaller effect on behavioral variation (partial η^2^ = 0.09, p < 0.001; Table 1). Predation regime also showed a statistically significant medium-sized interaction with species (partial η^2^ = 0.07, p < 0.001; Table 1). However, we found no evidence for a standard length * predation regime interaction (η^2^ = 0.04, p = 0.139; Table 1). Since the species did differ significantly in standard length (t = -14.368, p < 0.001), we examined whether the species * predation regime and species terms were masking a potential standard length * predation regime interaction effect in the MANCOVA. Even without these terms in the model, there was no sign of a standard length * predation regime effect (p = 0.29), suggesting that the interaction was driven by species identity rather than standard length.

**Table 1.** MANCOVA results table showing partial η**²**, F-value, and p-value derived from a Type III Wilks’ lambda multivariate test. Effects are organized by each explanatory variable in the model.

| **Response variables: Latency to barrier (Social latency) + Solve time (Inhibitory control) + Distance moved (Exploration) + Time in black thigmotaxis (Shyness)+ Time in white (Boldness)** |  |  |  |
| **Explanatory variable** | **Partial η²** | **F-value** | **p-value** |
|---|---|---|---|
| Predation Regime | **0.15** | **10.3** | **<0.001** |
| Species | **0.3** | **18.9** | **<0.001** |
| Replicate * Predation Regime | **0.09** | **6.4** | **<0.001** |
| Species * Predation Regime | **0.07** | **5.2** | **<0.001** |
| Standard Length * Predation Regime | 0.04 | 1.7 | 0.139 |

In our LDA examining categorization by species and predation regime across all individuals, we found that LD1 was primarily determined by social latency and, to a lesser extent, shyness (Figure 2A). In contrast, the primary loading contributing to LD2 was boldness.

Groups differed significantly in LD1 (Figure 2C, ANOVA; F = 72.26, p < 2e-16). *G. affinis* low-predation did not differ from high-predation *G. affinis* (Figure 2C). However, both differed from each of the *P. latipinna* groups, and the *P. latipinna* groups differed from one another (Figure 2C, Tukey’s HSD, p < 0.01). Groups also differed in LD2 (Figure 2D, ANOVA; F = 12.54, 8.05e-08). In LD2, low-predation *P. latipinna* had the highest LD2 score and differed significantly from each of the other groups, including the *P. latipinna* low-predation group (Figure 2D, p-adjusted < 0.05). Neither *G. affinis* group differed from one another, or from the high-predation *P. latipinna*, which had the lowest LD2 score (Figure 2D, p-adjusted > 0.05). Next, we cross-validated our four a priori categories using leave-one-out cross-validation (LOOCV). We found that the pre-defined categories were modestly robust, with a mean cross-validation accuracy of 45.5%, exceeding baseline expectations of 26.6% based on random permutation. The two high-predation categories were less separable than the low-predation categories. 43.40% of the observed high-predation *G. affinis* were correctly assigned to that category, and 40.24% of the high-predation *P. latipinna* were correctly assigned. In comparison, 51.81% of the observed low-predation *P. latipinna* and 48.45% of the low-predation *G. affinis* were correctly assigned to those categories. This level of clustering is consistent with the prior MANCOVA results.We did not find any evidence of sex-dependent effects of species or predation regime across any of the LMs constructed for the different response variables (Table S4). Thus, in the main text, we retained simpler LMs that included sex only as a covariate. In our LM, detour latency to engagement did not show general, species-specific, or size-specific effects of predation but did reveal a predation regime * replicate interaction (partial ^2^ = 0.03, F = 6.81, adjusted p = 0.003; Table 2; Figure 3). There was also a general species effect on latency to engagement (partial ^2^ = 0.10, F = 43.48, adjusted p < 0.001; Table 2). Specifically, *G. affinis* showed lower engagement latency than *P. latipinna* at each location (EMMs; p < 0.05; Figure 2A). Detour solve latency showed no evidence of a predation regime main effect or interactions with body size or species (Table 2; Figure 3).

**Table 2.** Linear model (LM) results for each response. P-values adjusted for multiple comparisons with false discovery rate correction based on the number of individual LMs constructed. Partial η² , F-values, and p-values were derived from Type III tests. “Replicate” here refers to the watershed identity (Replicate 1: Brushy Creek, Replicate 2: San Marcos River, or Replicate 3: Guadalupe River). Significant effects (p-value adjusted for false discovery rate < 0.05) are shown in bold.

| **Explanatory Variable** | **partial η<sup>2</sup>** | **F-value** | **p-value** |
|---|---|---|---|
| **Response: Detour time to engagement (Social motivation)** |  |  |  |
| Predation regime | <0.001 | 0.07 | 0.799 |
| Species | **0.10** | **43.48** | **<0.001** |
| Replicate * Predation regime | **0.03** | **6.81** | **0.003** |
| Species * Predation regime | 0.008 | 3.07 | 0.134 |
| Standard length * Predation regime | 0.001 | 0.41 | 0.652 |
| **Response: Detour solve time (Inhibitory control)** |  |  |  |
| Predation regime | <0.001 | 0.17 | 0.799 |
| Species | **0.01** | **4.01** | **0.049** |
| Replicate* Predation regime | <0.001 | 0.06 | 0.944 |
| Species * Predation regime | 0.003 | 1.08 | 0.374 |
| Standard length * Predation regime | 0.002 | 0.63 | 0.652 |
| **Response: Distance moved in scototaxis (Exploration)** |  |  |  |
| Predation regime | 0.008 | 2.93 | 0.147 |
| Species | **0.09** | **36.82** | **<0.001** |
| Replicate * Predation regime | 0.001 | 0.25 | 0.944 |
| Species * Predation regime | **0.02** | **6.28** | **0.032** |
| Standard length * Predation regime | <0.001 | 0.20 | 0.657 |
| **Response: Time in white zone (Boldness)** |  |  |  |
| Predation regime | **0.05** | **19.35** | **<0.001** |
| Species | **0.02** | **5.60** | **0.023** |
| Replicate * Predation regime | 0.02 | 3.11 | 0.077 |
| Species * Predation regime | **0.02** | **8.60** | **0.01** |
| Standard length * Predation regime | 0.006 | 2.27 | 0.332 |
| **Response: Time in black thigmotaxis (Shyness)** |  |  |  |
| Predation regime | **0.03** | **10.90** | **0.003** |
| Species | **0.03** | **12.17** | **<0.001** |
| Replicate * Predation regime | **0.12** | **23.20** | **<0.001** |
| Species * Predation regime | 0.002 | 0.70 | 0.403 |
| Standard length * Predation regime | **0.02** | **7.40** | **0.034** |

There was no evidence for a main effect of predation regime on total distance moved, but the effect of predation regime differed between species (partial ^2^ = 0.02, F = 6.28, adjusted p = 0.032; Table 2). Estimated marginal means indicated that total distance moved was lower in high-predation than low-predation *P. latipinna* (p = 0.004), whereas there was no evidence of a difference between predation regimes in *G. affinis* (p = 0.432; Figure 3). Total distance moved also decreased with standard length, and we found no evidence that this pattern was affected by predation regime (EMMs of trends; LP: -0.217, HP: -0.172; Table S2). *G. affinis* moved a greater distance than *P. latipinna*, however (EMMs contrasts; p < 0.01; Table 2; Figure 3).

Time in white (boldness) was affected by predation regime (partial ^2^ = 0.05, F = 19.35, adjusted p < 0.001; Table 2), with no evidence that the effect differed among replicates (partial ^2^ = 0.02, F = 3.11, adjusted p = 0.077; Table 2). High-predation fish spent more time in white than low-predation fish in two of the three replicates (Table S2). The effect of predation regime on time spent in white was also species-specific (partial ^2^ = 0.02, F = 8.60, adjusted p = 0.018; Table 2; Figure 4). Estimated marginal means indicated greater boldness in high-predation than low-predation *P. latipinna* across all three replicates (p < 0.001; Figure 4; Table S2), whereas there was no evidence of a difference between predation regimes in *G. affinis* (p = 0.448; Figure 4; Table S2).

**Figure 4.**
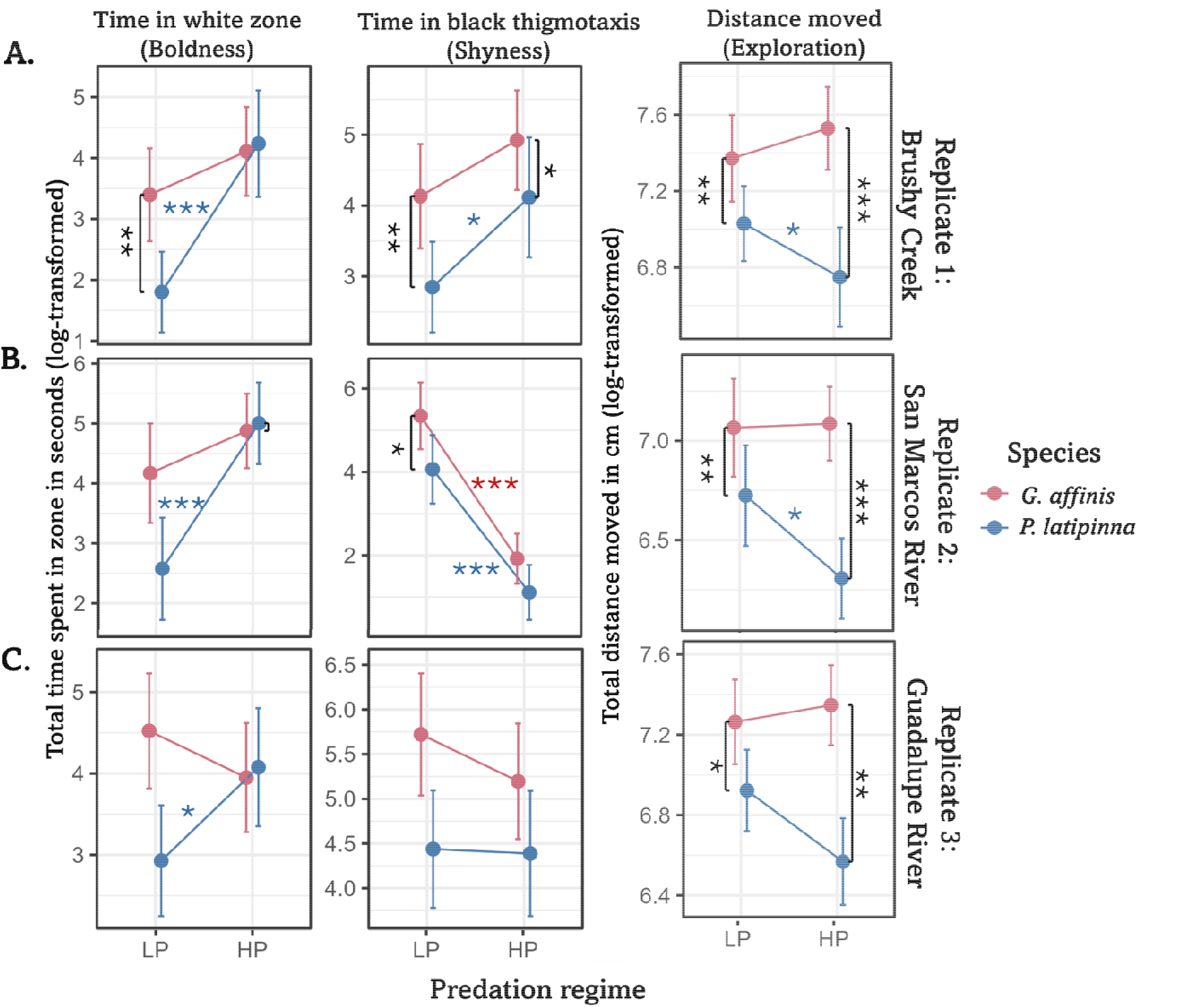
Estimates of boldness, shyness, and exploration measures by species and predation regime in (A.) Replicate (Brushy Creek), (B.) Replicate 2 (San Marcos River), and (C) Replicate 3 (Guadalupe River). Vertical asterisks and square brackets indicate significant species differences in model-adjusted estimated marginal means (EMMs) at a particular location. Horizontal asterisks are color-coded by species and indicate significant differences in EMMs between predation regimes within a watershed. For a given contrast, one asterisk indicates a p-value < 0.05, two indicates a p-value < 0.01, and three indicates a p-value < 0.001. EMMs for detecting species differences are derived from linear models (LMs) that includes a three-way interaction between species, predation regime, and watershed, in addition to sex and standard length covariates. Estimated marginal means for predation regime differences use the same LMs employed elsewhere in the study (see Methods).

Time in black thigmotaxis (shyness) differed between high- and low-predation populations (partial η^2^ = 0.03, F = 10.90, p = 0.003; Table 2), but this difference varied strongly among replicates (partial η^2^ = 0.12, F = 23.20, adjusted p < 0.001; Table 2). Thus, unlike in time in white, there was little evidence for a consistent association between predation regime and time in black thigmotaxis across replicate watersheds. The difference between predation regimes also varied with standard length (partial η^2^ = 0.02, F = 7.40, adjusted p = 0.034; Table 2).

## Discussion

We predicted that predation-associated divergence would be (1) consistent across replicated predation regimes, (2) stronger in the species with more conspicuous mating behavior and morphology (*Poecilia latipinna*), and/or (3) stronger in larger-bodied individuals. At the multivariate level, we found that predation-associated divergence was primarily species-specific rather than shared with minimal influence of body size (Table 1, Figure 2). Furthermore, the species-specific comparisons suggest that specific behavioral traits emerge as a possible interaction between sexual selection and natural selection, with divergence magnified (e.g. social motivation, Figure 2C) or minimized (e.g. boldness, Figure 2D) between species with different male mating tactics and morphology under greater predation risk. Multivariate divergence in the social motivation-exploration axis was stronger between predation regimes in *P. latipinna* than between the two species, supporting the notion that predation may be associated with different behavioral optima across multiple measures of behavior and cognition for the conspicuous prey species, but not the less conspicuous one (Figure 2C). Meanwhile, species-specific differences observed under the low-predation regime in the boldness multivariate axis were absent in the high-predation regime (Fig 2D), suggesting that predation pressure has a stronger influence than sexual selection on specific behavioral traits.

Interestingly, some behaviors were less sensitive to predation risk but exhibited consistent species-level differences across predation regimes. Social motivation consistently differed between the species across replicates independently predation regime (Table 2; Figure 3). This is consistent with predictions that sexual selection strongly shapes social interactions between species with different mating systems but differs from previous findings that demonstrate an effect of predation on social aggregation (e.g. in Trinidadian guppies Seghers 1974; Magurran and Seghers 1990). Despite the prevalence of shoaling in high-predation guppies and its potential to function as a dilution mechanism, shoaling might also increase detectability to predators, and thus it might not be universally favored in environments with higher predator abundance. The strong species difference we observed in social motivation (Table 2; Figure 3) is consistent with the possibility that male mating tactics can shape behavior independently of predation. The exclusively coercive species *G. affinis* showed consistently greater social motivation than the mixed-tactic *P. latipinna*. This finding is consistent with previous studies comparing female social aggregation tendencies of mixed-tactic and all-coercive poeciliid fish species (Dadda et al. 2005; Dadda 2015; Inman et al. 2025a). *G. affinis* males exhibit high coercive mating rates and pursue females regardless of their receptivity (Deaton 2008). Thus, high social motivation in female *G. affinis* may dilute the adverse effects of being an individual target for coercive male conspecifics, while high social motivation in male *G. affinis* may favor greater coercion success.

At the univariate level, support for our predictions for predation-associated divergence was most evident in boldness and, to a lesser extent, exploration (Table 2; Figure 4). In contrast, associations between predation regime and shyness varied among replicates, providing evidence against a general predation regime or size-dependent response. Shyness also showed body size-dependent differences between predation regimes. However, the strong interaction with replicate affecting shyness more generally limits interpretation of this pattern. Generally, the effects of shared predation regimes detected at the multivariate level were reflected most clearly by boldness at the univariate level, with more limited support from exploration.

High-predation environments were generally associated with higher boldness (Figure 4, Table 2). This is consistent with a growing theoretical and empirical literature suggesting that a high baseline risk of predation may favor higher, rather than lower, boldness (Brown et al. 2005; Brown et al. 2007a; Fuiman et al. 2010; Harris et al. 2010; Blake et al. 2018). Animals that live under constant predation risk still must mate and acquire resources while also successfully evading detection or capture (Lima and Bednekoff 1999) and may thus appear relatively bold (Brown et al. 2005). However, these findings contrast with other work showing the opposite pattern of lower boldness in higher-predation sticklebacks (Brydges et al. 2008), *Brachyrhaphis rhabdophora* (Rasmussen and Belk 2017), and female *Gambusia hubbsi* (Heinen-Kay et al. 2016). These contrasting patterns may be due to multiple factors, including an organism’s foraging ecology, habitat structure, and whether predators represent a sporadic or constant threat in the environment (Lima 1987; Savino and Stein 1989; Lima and Bednekoff 1999).

Consistent with our second prediction, differences in boldness between predation regimes were species-dependent, with larger differences between high- and low-predation populations in the species with greater behavioral and morphological conspicuousness ( *P. latipinna*,Figure 4; Table 2; Table S2). *P. latipinna* engages in highly conspicuous courtship displays whereas *G. affinis* does not. *P. latipinna* courting males have blue and orange nuptial coloration and enlarged dorsal fins, often displaying prominently to females in well-lit areas during the breeding season (Ptacek and Travis 1996).As with ornaments in swordtails (Rosenthal et al. 2001; Basolo and Wagner 2004; Hernandez-Jiménez and Ríos-Cardenas 2012), conspicuous *P. latipinna* male morphology and behavior in the wild may allow predators to more effectively target *P. latipinna* individuals of both sexes than *G. affinis*. Greater vulnerability to predation may lead to higher encounter rates between predators and *P. latipinna* than between predators and *G. affinis*, favoring an increase in bold behavior in the former species under higher predation risk. Higher boldness may be favored in *P. latipinna* from high-predation environments because they must efficiently carry out routine activities like foraging and mating outside of an immediate predation threat (Lima and Bednekoff 1999). In contrast, populations of *Gambusia affinis*, may not experience different levels of vulnerability between the high- and low-predation regimes in our study if they are less conspicuous or preferred as prey, resulting in no consistent differences in boldness between predation regimes. Thus, reproductive phenotypes associated with sexual selection could contribute to the stronger behavioral divergence between predation regimes observed in *P. latipinna* than *G. affinis*.

While boldness and shyness responses were correlated, shyness showed substantially greater heterogeneity among replicate watersheds than boldness (Figure 4; Table 2). The differences between predation regimes also varied with standard length, but the strong heterogeneity among watersheds limits interpretation of this size-dependent pattern as a general response to predation. Other ecological factors, such as light environment, structural complexity, or other differences among populations, may contribute to variation in shyness.

Prior theoretical and empirical work suggests that a reduction in conspicuous movements might lower predation risk (Werner and Anholt 1993; Reale et al. 2007; Richardson 2001). In keeping with these predictions, we found an effect of predation regime on exploration in one of the two species we examined (Table 2, Figure 4). Specifically, the more conspicuous *P. latipinna* showed directionally lower exploration in all three watersheds (Table 2; Figure 4). This implies that *P. latipinna* might compensate for increased boldness with reduced movement under threatening conditions. However, the species effect on exploration was much larger than any other factor, with *G. affinis* showing higher exploration than *P. latipinna* across 6 locations (Table 2; Figure 4).

The species difference observed in exploration is not consistent with findings in predator-free conditions, in which females of species with exclusively coercive male mating tactics showed consistently lower exploration than species with mixed-tactic mating systems (Inman et al. 2025a). However, such patterns did not hold when additional species were compared, including *G. affinis* and *P. latipinna* (Inman et al. 2025b). peasons Foraging ecology is also another possible for the species difference we observed. *G. affinis* is a generalist omnivore, while *P*. *latipinna* has a more specialized algivorous diet (Harrington and Harrington 1961; Lee et al. 2018; Pyke 2005; Alkahem et al. 2007), potentially favoring greater exploratory movement in *G. affinis*.

Although these results suggest that species’ traits may shape behavioral divergence between predation regimes, identifying mechanisms underlying this asymmetry will require more direct study of predator-prey interactions and their context. For example, future work in this system or others with similar sympatric species pairs could directly test our conjectures about relative predation vulnerability by directly measuring mortality rates (e.g. Krebs and Bell 2012; Johnson and Zúñiga-Vega 2009) or predator attack rates on prey. This study was also unable to disentangle experiential effects of predation risk from heritable effects (Brown et al. 2007a), or consumptive from nonconsumptive effects (Preisser and Bolnick 2008). Another important consideration is diversity in predator species: predators may vary in important ways in their feeding strategies and preferences (Carey and Wahl 2011; Falk et al. 2015). In referring to locations as “high-predation,” this study generalized across a range of fish predator species from centrarchids to cichlids. Studies of predator-prey interactions across multiple predator species in experimental mesocosms can yield insights into phenotype-specific predation risk in nature (e.g. Richardson et al. 2006).

In summary, we show that comparing sympatric species across shared environments can reveal variation in behaviors associated with predation regime that would be missed by studying either species alone. More broadly, considering species differences in reproductive phenotypes associated with sexual selection and other traits shape predation vulnerability may improve predictions about when predation regimes generate shared or species-specific behavioral divergence.

## Supporting information

Supplementary Methods and Results

## Acknowledgements

We thank Marina Hutchins, Jojo Koley, Maxximus Ramsaroop, Alan Vuong, Mahati Sudhagar, Caylee Campbell, Max Roemer, Lara Maschio, Gita Kolluru, Ingo Schlupp, and Phil Queller for their assistance in the field. We are grateful to Aidyn Neiman, Micah McDowell, Isabel Ebben, Maddie Woodul, Catia Rodriguez, and Mary Ramsey for providing invaluable assistance in the lab. We also thank Carlos Botero, Larry Gilbert, Mike Ryan, Justin Yeager, Dan Powell, Marina Hutchins, Sunishka Thakur, Maxximus Ramsaroop, Savvy Cornett, Brittany Hamauei, and Sydney Jansen for providing critical feedback on earlier versions of this manuscript. UT Austin IACUC approved laboratory housing and field protocols for this project (Protocols AUP-2022-00224 and AUP-2025-2200224). CMI thanks the Texas Parks & Wildlife Department for both a Scientific Research Permit (SPR-0725-059) and a Texas Fishing License to complete this work, as well as Braun & Gresham, PLLC’s Texas Ecological Laboratories program for funding. MEC thanks NSF for funding (IOS-1911826 award to MEC).

