## Supplementary Methods and Results for "Behavioral divergence across replicated predation regimes is species-specific in two sympatric poeciliid fishes"

**Supplementary Material**

*Methods (Supplemental)*

We constructed a supplemental MANCOVA model with a “Sex * Species * Predation regime” interaction term due to the often sex-specific nature of predation effects on behavior in similar systems, such as the Trinidadian guppy (Godin 1995; Brusseau et al. 2023). We also constructed two supplemental LMs: one was constructed to detect potential sex-dependent effects and the other was to detect species differences within locations for purposes of visualization. The first included a three-way “Sex * Species * Predation regime” interaction in addition to terms included in the other models. Though we did not develop a priori hypotheses about the magnitude and direction of species differences, we constructed a separate LM for each behavior with a three-way interaction between watershed, predation regime, and species (rather than separate two-way interactions) that allowed species contrasts to be computed at each location (vertical brackets and asterisks shown in Figures 3-4) but was otherwise the same in structure as the original set of LMs. Because the primary aim of these visualizations was to show species-specific predation regime effects rather than species differences, the estimated marginal means depicted in the figures are derived from the LMs used for hypothesis testing.

*Results (Supplemental)*

The MANCOVA that included a three-way interaction between sex, species, and predation regime provided little evidence that sex modified species-specific responses to predation (Table S3). Because sex-specific responses were not among our a priori hypotheses, we retained sex as a main-effect covariate in the primary models and report the expanded model here in Supplementary Material (Table S3). In the LMs, none of the interactions involving sex were supported across all five traits examined (Table S4). Linear models conducted on each behavioral and cognitive response variable used the same set of explanatory variables employed in the earlier MANCOVA. To account for possible Type I errors across multiple LMs, P-values were adjusted using a false discovery rate correction (Benjamini and Hochberg 1995).

**Table S1.** Estimated marginal means for general, species-specific, and size-specific predation regime effects. Estimated marginal means contrasts are derived from the mixed effects ANCOVA model constructed for each response variable.

| **Explanatory variable** | **Contrast** | **Contrast estimate** | t**.ratio** | **p-value** |
| --- | --- | --- | --- | --- |
| **Response: Detour latency to engagement** | | | |  |
| Predation regime | LP- HP | -0.022 | -0.21 | 0.836 |
| Predation regime * Replicate | Brushy Creek: LP-HP | 0.545 | 2.62 | 0.009 |
|  | San Marcos R: LP-HP | -0.421 | -2.52 | 0.012 |
|  | Guadalupe R: LP-HP | -0.190 | -1.09 | 0.278 |
| Predation regime * Species | *G. affinis*: LP-HP | 0.198 | 1.27 | 0.205 |
|  | *P. latipinna*: LP-HP | -0.242 | -1.39 | 0.165 |
| Predation regime * Standard length | LP-HP (slope difference) | -0.092 | -0.64 | 0.522 |
| **Response: Latency to solve** | | | | |
| Predation regime | LP- HP | 0.037 | 0.339 | 0.735 |
| Predation regime * Replicate | Brushy Creek: LP-HP | 0.072 | 0.343 | 0.731 |
|  | San Marcos R: LP-HP | -0.013 | -0.07 | 0.941 |
|  | Guadalupe R: LP-HP | 0.051 | 0.29 | 0.773 |
| Predation regime * Species | *G. affinis*: LP-HP | -0.094 | -0.608 | 0.544 |
|  | *P. latipinna*: LP-HP | 0.168 | 0.952 | 0.342 |
| Predation regime * Standard length | LP-HP (slope difference) | 0.117 | 0.796 | 0.427 |
| **Response: Total distance moved** | | | | |
| Predation regime | LP- HP | 0.132 | 1.72 | 0.087 |
| Predation regime * Replicate | Brushy Creek: LP-HP | 0.062 | 0.45 | 0.655 |
|  | San Marcos R: LP-HP | 0.199 | 1.45 | 0.147 |
|  | Guadalupe R: LP-HP | 0.136 | 1.12 | 0.263 |
| Predation regime * Species | *G. affinis*: LP-HP | -0.087 | -0.79 | 0.432 |
|  | *P. latipinna*: LP-HP | 0.351 | 2.87 | 0.004 |
| Predation regime * Standard length | LP-HP (slope difference) | -0.044 | -0.44 | 0.657 |
| **Response: Time in white** | | | | |
| Predation regime | LP- HP | **-1.14** | **-4.41** | **<0.001** |
| Predation regime * Replicate | Brushy Creek: LP-HP | **-1.574** | **-3.38** | **<0.001** |
|  | San Marcos R: LP-HP | **-1.567** | **-3.41** | **<0.001** |
|  | Guadalupe R: LP-HP | -0.288 | -0.71 | 0.479 |
| Predation regime * Species | *G. affinis*: LP-HP | -0.282 | -0.76 | 0.448 |
|  | *P. latipinna*: LP-HP | **-2.004** | **-4.88** | **<0.001** |
| Predation regime * Standard length | LP-HP (slope difference) | 0.504 | 1.51 | 0.133 |
| **Response: Shyness** | | | | |
| Predation regime | Low Predation- High Predation | 0.817 | 3.26 | 0.001 |
| Predation regime * Replicate | Brushy Creek: LP-HP | -1.029 | -2.29 | 0.02 |
|  | San Marcos R: LP-HP | 3.191 | 7.18 | <0.001 |
|  | Guadalupe R: LP-HP | 0.287 | 0.731 | 0.465 |
| Predation regime * Species | *G. affinis*: LP-HP | 1.054 | 2.94 | 0.004 |
|  | *P. latipinna*: LP-HP | 0.579 | 0.29 | 0.772 |
| Predation regime * Standard length | LP-HP (slope difference) | **0.881** | **2.721** | **0.007** |

| Response | PC1 (51.35%) | PC2 (25.81%) | PC3 (22.83%) |
| --- | --- | --- | --- |
| Shyness | -0.604 | 0.193 | 0.774 |
| Boldness | -0.548 | -0.805 | -0.226 |
| Exploration | -0.579 | 0.561 | -0.592 |

**Table S2.** PC loadings for time in black thigmotaxis (shyness), time in white (boldness), and total distance moved (exploration). The variance explained by each PC is included in parentheses.

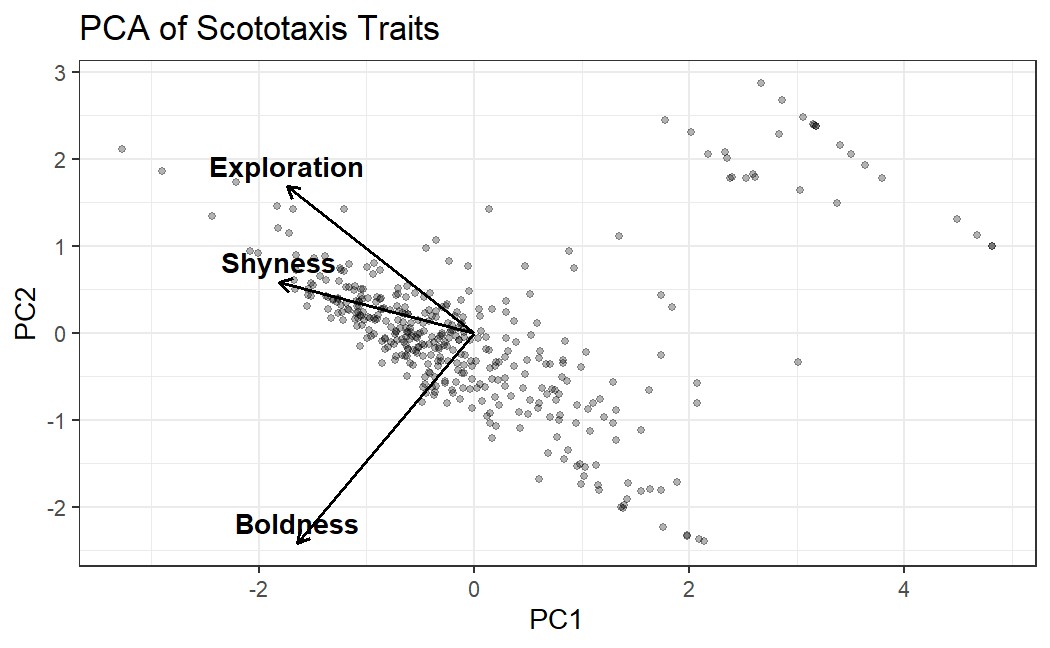

**Figure S1.** Biplot illustrating Principal Component loadings of total distance moved (exploration), time in black thigmotaxis (shyness), and time in white (boldness). Exploration, shyness, and boldness are directionally similar in PC1. PC1 explained 51.35% of the variance among the three factors, while PC2 explained 25.81% of the variance.

**Table S3**. MANCOVA results for model that includes three-way interaction between sex, predation regime, and species. Table shows partial η**²**, F-value, and p-value derived from a Type III Wilks’ lambda multivariate test. Effects are organized by each explanatory variable in the model. Since the interactions between sex, species, and predation regime were not significant (and did not change other patterns), sex was excluded from the interaction in the MANCOVA model presented in the main text.

| **Response variables: Latency to barrier (Social latency) + Solve time (Inhibitory control) + Distance moved (Exploration) + Time in black thigmo (Shyness)+ Time in white (Boldness)** | | | |
| --- | --- | --- | --- |
| **Explanatory variable Partial η² F-value p-value** | | | |
| Predation regime | **0.15** | **2.30** | **0.045** |
| Species | **0.24** | **15.90** | **<0.001** |
| Replicate * Predation regime | **0.09** | **6.14** | **<0.001** |
| Species * Predation regime | **0.06** | **4.49** | **<0.001** |
| Sex * Predation regime | 0.06 | 1.56 | 0.172 |
| Sex * Species * Predation regime | 0.02 | 1.34 | 0.248 |
| Standard length * Predation regime | 0.04 | 0.993 | 0.422 |

**Table S4.** Results for each behavioral measure from LMs that included the three way “Sex * Species * Predation regime” interaction. P-values adjusted for multiple comparisons with false discovery rate correction based on the number of individual LMs constructed. All three variables are derived from a Type III ANCOVA test. “Replicate” here refers to the watershed identity (Replicate 1: Brushy Creek, Replicate 2: San Marcos River, or Replicate 3: Guadalupe River).

| **Explanatory Variable** | **partial η²** | **F-value** | **p-value** |
| --- | --- | --- | --- |
| **Response: Detour time to engagement (Social motivation)** | | | |
| Predation regime | <0.001 | 0.14 | 0.704 |
| Species | **0.09** | **40.41** | **<0.001** |
| Replicate * Predation regime | **0.03** | **6.90** | **0.003** |
| Species * Predation regime | 0.005 | 2.27 | 0.221 |
| Standard length * Predation regime | 0.001 | 0.530 | 0.584 |
| Sex * Species * Predation regime | 0.002 | 0.684 | 0.681 |
| Sex * Predation regime | <0.001 | 0.02 | 0.892 |
| **Response: Detour solve time (Inhibitory control)** | | | |
| Predation regime | <0.001 | 0.36 | 0.689 |
| Species | 0.008 | 3.53 | 0.061 |
| Replicate * Predation regime | 0.001 | 0.23 | 0.798 |
| Species * Predation regime | <0.001 | 0.08 | 0.783 |
| Standard length * Predation regime | 0.006 | 2.68 | 0.256 |
| Sex * Species * Predation regime | <0.001 | 0.20 | 0.817 |
| Sex * Predation regime | 0.008 | 3.29 | 0.088 |
| **Response: Distance moved in scototaxis (Exploration)** | | | |
| Predation regime | 0.007 | 2.69 | 0.169 |
| Species | **0.08** | **33.69** | **<0.001** |
| Replicate * Predation regime | 0.002 | 0.31 | 0.798 |
| Species * Predation regime | 0.02 | 6.23 | 0.065 |
| Standard length * Predation regime | <0.001 | 0.59 | 0.584 |
| Sex * Species * Predation regime | 0.005 | 1.78 | 0.678 |
| Sex * Predation regime | <0.001 | 1.23 | 0.381 |
| **Response: Time in white zone (Boldness)** | | | |
| Predation regime | **0.06** | **22.20** | **<0.001** |
| Species | **0.02** | **5.641** | **0.023** |
| Replicate * Predation regime | 0.02 | 2.98 | 0.086 |
| Species * Predation regime | 0.01 | 4.49 | 0.087 |
| Standard length * Predation regime | <0.001 | 0.16 | 0.691 |
| Sex * Species * Predation regime | <0.001 | 0.003 | 0.955 |
| Sex * Predation regime | 0.01 | 3.79 | 0.176 |
| **Response: Time in black thigmotaxis (Shyness)** | | | |
| Predation regime | **0.03** | **10.42** | **0.003** |
| Species | **0.03** | **12.51** | **<0.001** |
| Replicate * Predation regime | **0.12** | **23.75** | **<0.001** |
| Species * Predation regime | 0.002 | 0.43 | 0.643 |
| Standard length * Predation regime | 0.01 | 4.17 | 0.209 |
| Sex * Species * Predation regime | 0.003 | 1.22 | 0.678 |
| Sex * Predation regime | 0.003 | 1.06 | 0.381 |

**Table S5.** Repeatability estimates in the scototaxis assay across all individuals, each species, each predation regime, or species-predation regime combinations. Repeatability was derived from a subset of individuals (n = 101) across both species collected from two locations: high-predation Replicate 1 and low-predation Replicate 1 (Brushy Creek watershed). Repeatability was estimated using intra-class correlation coefficients (written as “R”), with D and p-values derived from likelihood ratio tests on linear mixed models (LMMs). See Methods in main text for model details.

| **Group** | **R ± SE** | **D** | **p-value** |
| --- | --- | --- | --- |
| **Time in white (Boldness)** | | | |
| Overall | 0.39 ± 0.08 | 16.5 | <0.001 |
| *P. latipinna* | 0.31 ± 0.13 | 4.77 | 0.01 |
| *G. affinis* | 0.35 ± 0.12 | 7.15 | 0.003 |
| High-Predation | 0.61 ± 0.11 | 14.2 | <0.001 |
| Low-Predation | 0.32 ± 0.11 | 7.85 | 0.003 |
| *P. latipinna* HP | 0.60 ± 0.22 | 4.53 | 0.02 |
| *G. affinis* HP | 0.25 ± 0.19 | 1.31 | 0.13 |
| *P. latipinna* LP | 0.25 ± 0.15 | 2.53 | 0.06 |
| *G. affinis* LP | 0.36 ± 0.15 | 4.69 | 0.02 |
| **Time in black thigmotaxis** | | | |
| Overall | 0.35 ± 0.09 | 13.4 | <0.001 |
| *P. latipinna* | 0.25 ± 0.13 | 3 | 0.04 |
| *G. affinis* | 0.48 ± 0.11 | 13.6 | <0.001 |
| High-Predation | 0.18 ± 0.15 | 0.92 | 0.17 |
| Low-Predation | 0.35 ± 0.1 | 9.53 | 0.001 |
| *P. latipinna* HP | 0 ± 0.18 | 0 | 1 |
| *G. affinis* HP | 0.89 ± 0.05 | 31.4 | <0.001 |
| *P. latipinna* LP | 0.31 ± 0.14 | 3.77 | 0.03 |
| *G. affinis* LP | 0.42 ± 0.14 | 6.48 | 0.005 |
| **Distance moved in scototaxis (Exploration)** | | | |
| Overall | 0.62 ± 0.06 | 48.7 | <0.001 |
| *P. latipinna* | 0.61 ± 0.09 | 22 | <0.001 |
| *G. affinis* | 0.34 ± 0.12 | 6.38 | 0.005 |
| High-Predation | 0.72 ± 0.10 | 21.8 | <0.001 |
| Low-Predation | 0.60 ± 0.08 | 30.9 | <0.001 |
| *P. latipinna* HP | 0.83 ± 0.13 | 11.5 | <0.001 |
| *G. affinis* HP | 0.47 ± 0.18 | 5.02 | 0.01 |
| *P. latipinna* LP | 0.56 ± 0.11 | 14.3 | <0.001 |
| *G. affinis* LP | 0.23 ± 0.15 | 1.86 | 0.09 |

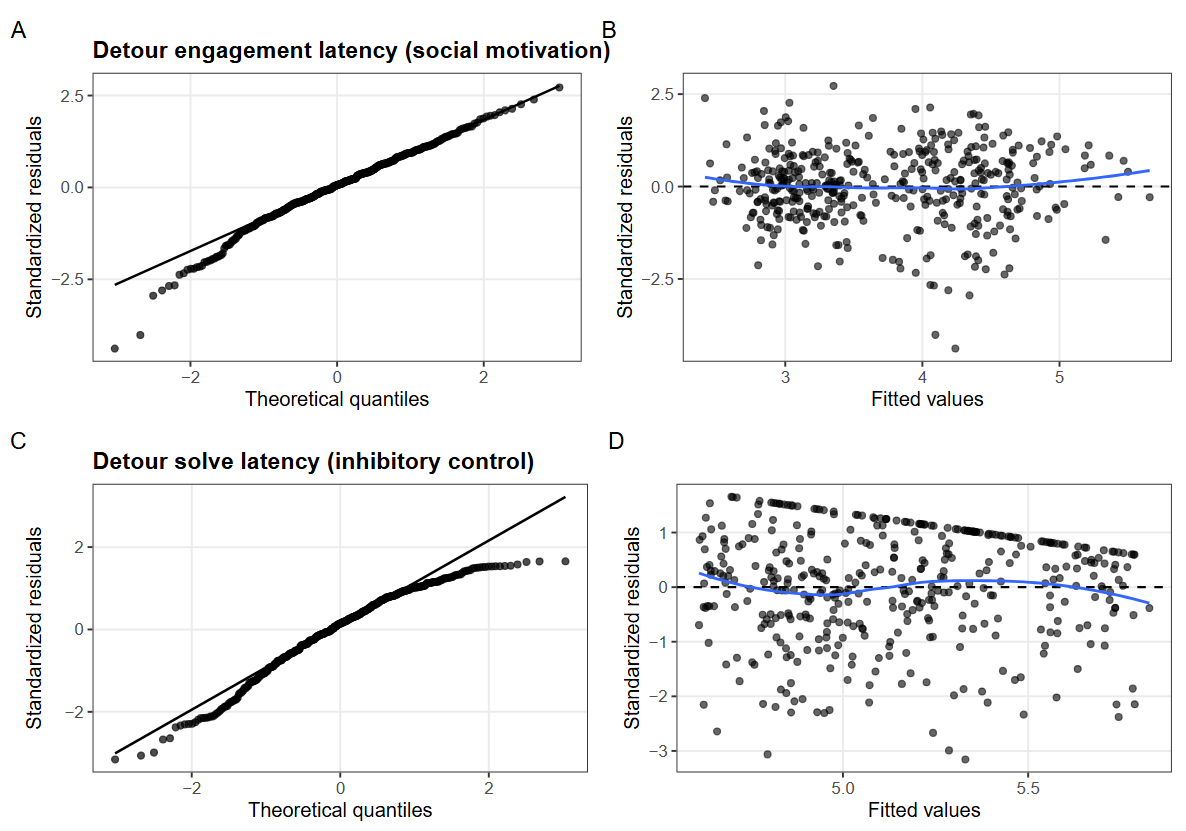
**Figure S2.** Diagnostics of the two measures taken in the detour reaching assay. Panel A shows the Quantile-Quantile plot of standardized against theoretical residuals for detour engagement latency (social motivation), while Panel B shows standardized residuals plotted against model-predicted fitted values for the same measure. Panel C and D are the same plots applied to detour solve latency (inhibitory control).

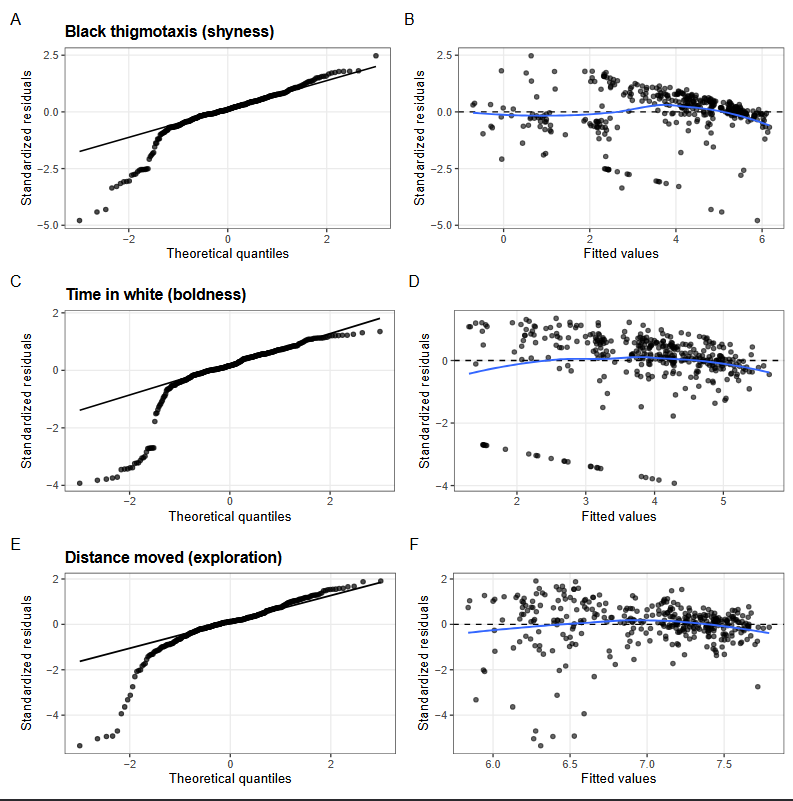

**Figure S3.** Diagnostics of the three measures taken in the scototaxis assay. Panel A shows the Quantile-Quantile plot of standardized against theoretical residuals for time in black thigmotaxis (shyness), while Panel B shows standardized residuals plotted against model-predicted fitted values for the same measure. Panel C and D are the same plots applied to time in white (boldness), while E and F show plots for distance moved (exploration).

**Table S6.** Diagnostic tests for linear models of behavioral responses. P-values are reported for DHARMa simulation-based tests of residual uniformity, dispersion, and outliers (1000 simulations per model). The number and percentage of observations with absolute standardized residuals > 3 are also reported. N indicates the number of observations in each model.

| **Response** | **N** | **DHARMa uniformity p-value** | **Dispersion p-value** | **Outlier p-value** | **\|standardized residual\| > 3, n (%)** |
| --- | --- | --- | --- | --- | --- |
| Time in black thigmotaxis (shyness) | 364 | <0.001 | 0.698 | <0.001 | 9 (2.5%) |
| Time in white (boldness) | 364 | <0.001 | 0.698 | <0.001 | 17 (4.7%) |
| Distance moved (exploration) | 364 | <0.001 | 0.694 | <0.001 | 2 (0.5%) |
| Detour engagement latency (Social motivation) | 412 | 0.312 | 0.694 | 0.199 | 2 (0.5%) |
| Detour solve latency (Inhibitory control)4 | 412 | 0.107 | 0.694 | 1 | 2 (0.5%) |

**Table S7**. Comparison of primary and sensitivity analyses of individual response variables across effects of interest. Sensitivity analyses were performed by running a linear model on each response variable with the outliers (total number in column ‘N_excluded’ that were identified by DHARMa diagnostics excluded (Table S6). P-values presented are from Type III F-tests performed on the linear model.

| **Response** | **Effect** | **N_excluded** | **p_primary** | **p_sensitivity** |
| --- | --- | --- | --- | --- |
| Time in black thigmotaxis (shyness) | Predation regime | 9 | 0.001 | <0.001 |
|  | Predation regime * Species | 9 | 0.403 | 0.098 |
|  | Predation regime * Standard length | 9 | 0.007 | 0.003 |
|  | Predation regime * Replicate | 9 | <0.001 | <0.001 |
| Time in white (boldness) | Predation regime | 17 | <0.001 | <0.001 |
|  | Predation regime * Species | 17 | 0.004 | <0.001 |
|  | Predation regime * Standard length | 17 | 0.133 | 0.094 |
|  | Predation regime * Replicate | 17 | 0.046 | 0.115 |
| Total distance moved (exploration) | Predation regime | 9 | 0.088 | 0.012 |
|  | Predation regime * Species | 9 | 0.013 | 0.009 |
|  | Predation regime * Standard length | 9 | 0.657 | 0.708 |
|  | Predation regime * Replicate | 9 | 0.782 | 0.153 |
| Detour engagement latency (Social motivation) | Predation regime | 2 | 0.799 | 0.875 |
|  | Predation regime * Species | 2 | 0.081 | 0.152 |
|  | Predation regime * Standard length | 2 | 0.522 | 0.483 |
|  | Predation regime * Replicate | 2 | 0.001 | <0.001 |
| Detour solve latency (inhibitory control) | Predation regime | 2 | 0.678 | 0.491 |
|  | Predation regime * Species | 2 | 0.299 | 0.206 |
|  | Predation regime * Standard length | 2 | 0.427 | 0.597 |
|  | Predation regime * Replicate | 2 | 0.944 | 0.894 |
